# Stationary Ca^2+^ nanodomains in the presence of buffers with two binding sites

**DOI:** 10.1101/2020.09.14.296582

**Authors:** Y. Chen, V. Matveev

## Abstract

We examine closed-form approximations for the equilibrium Ca^2+^ concentration near a point Ca^2+^ source representing a Ca^2+^ channel, in the presence of a mobile Ca^2+^ buffer with 2:1 Ca^2+^ binding stoichiometry. We consider buffers with two Ca^2+^ binding sites activated in tandem and possessing distinct binding affinities and kinetics. This allows to model the impact on Ca^2+^ nanodomains of realistic endogenous Ca^2+^ buffers characterized by cooperative Ca^2+^ binding, such as calretinin. The approximations we present involve a combination or rational and exponential functions, whose parameters are constrained using the series interpolation method that we recently introduced for the case of 1:1 Ca^2+^ buffers. We conduct extensive parameter sensitivity analysis and show that the obtained closed-form approximations achieve reasonable qualitative accuracy for a wide range of buffer’s Ca^2+^ binding properties and other relevant model parameters. In particular, the accuracy of the newly derived approximants exceeds that of the rapid buffering approximation in large portions of the relevant parameter space.

**STATEMENT OF SIGNIFICANCE:** Closed-form approximations describing equilibrium distribution of Ca^2+^ in the vicinity of an open Ca^2+^ channel proved useful for the modeling of local Ca^2+^ signals underlying secretory vesicle exocytosis, muscle contraction and other cell processes. Such approximations provide an efficient method for estimating Ca^2+^ and buffer concentrations without computationally expensive numerical simulations. However, while most biological buffers have multiple Ca^2+^ binding sites, much of prior modeling work considered Ca^2+^ dynamics in the presence of Ca^2+^ buffers with a single Ca^2+^ binding site. Here we extend modeling work on equilibrium Ca^2+^ nanodomains to the case of Ca^2+^ buffers with two binding sites, allowing to gain deeper insight into the impact of more realistic Ca^2+^ buffers, including cooperative buffers, on cell Ca^2+^ dynamics.

## INTRODUCTION

Accurate description of Ca^2+^ concentration ([Ca^2+^]) elevations formed near open Ca^2+^ channels, termed micro- or nano-domains, is crucial for the understanding of many fundamental cell processes such as synaptic neurotransmitter release, endocrine hormone release, and muscle contraction (1-5). This is particularly true in the case of chemical synaptic transmission, since the fusion (exocytosis) of a presynaptic neurotransmitter-filled vesicle can be triggered by the opening of just a few voltage-gated Ca^2+^ channels (5-10). The characteristic time of synaptic vesicle exocytosis is a fraction of 1 millisecond, while the relevant spatial scale is determined by the Ca^2+^ channel-vesicle separation, on the order of 10-100 nm (4, 5, 10-15). Optical Ca^2+^ imaging is insufficient to track spatio-temporal Ca^2+^ dynamics on such fine temporal and spatial scales, and cannot be carried out without disturbing the Ca^2+^ signal that is being measured. This explains the key role that mathematical and computational modeling has played in the study of vesicle exocytosis, myocyte contraction, and other fundamental processes controlled by localized Ca^2^ elevations (13-20). The main technical challenge in such modeling stems from the interaction of Ca^2+^ with intracellular Ca^2+^ buffers, which bind most of Ca^2+^ ions upon their entry into the cytoplasm (12, 19). Buffered Ca^2+^ diffusion problem leads to a system of nonlinear partial differential equations, which requires computational modeling. One early insight gained from such computational studies is that [Ca^2+^] reaches a quasi-stationary steady state in the vicinity of an open Ca^2+^ channel very quickly, within 10-100 μs, and this quasi-stationary Ca^2+^ nanodomain gradient collapses as quickly after channel closing (15-20). This suggested that equilibrium solutions to the Ca^2+^ reaction-diffusion equations achieve sufficient accuracy in modeling [Ca^2+^] as a function of distance from an open Ca^2+^ channel. Therefore, several closed-form equilibrium Ca^2+^ nanodomain approximations have been developed, most notably the excess buffer approximation (EBA), the rapid buffering approximation (RBA) and the linear approximation (LIN) (19, 21-31). These approximations proved useful in understanding the properties of Ca^2+^ nanodomains and their dependence on the properties of cell Ca^2+^ buffers, and provide a convenient and efficient tool for modeling studies (19, 26, 32-36). More recently, we introduced two new methods for finding such analytic approximations, one of which is based on the variational approach, and the second method based on matching the coefficients of short-range Taylor series and long-range asymptotic series of the nanodomain [Ca^2+^] as a function of distance from the channel (37, 38). Here we present an extension of the latter approach, which we refer to as the series interpolation method, to the case of more complex buffers with two Ca^2+^ binding sites. This allows to model the impact of more realistic Ca^2+^ buffers, all of which have multiple binding sites. For example, many widely expressed Ca^2+^ buffers and sensors such as calretinin and calmodulin contain two EF-hand domains which cooperatively bind two Ca^2+^ ions, whereby the binding of the second Ca^2+^ ion proceeds with much greater affinity once the first binding site is occupied (39-47). To date, only RBA has been extended to such realistic buffers, and it achieves sufficient accuracy only in restrictive parameter regimes corresponding to a very small ratio between the rates of diffusion and Ca^2+^ binding reactions (48). In this study we show that the series interpolation methods can be successfully extended to such buffers with more realistic Ca^2+^ binding properties, using simple *ansätze* combining exponential and rational functions, similar to those considered in (37) for the case of 1:1 buffers. We perform systematic parameter sensitivity analysis of the accuracy of the newly obtained approximants and demonstrate that they achieve significantly improved approximation accuracy as compared to RBA for a wide range of relevant parameter values, and capture the non-trivial dependence of the bound buffer concentration on the distance from the Ca^2+^ channel.

## METHODS

We start with the description of the Ca^2+^ binding and unbinding reactions for buffer molecules with two binding sites, which we will refer to as two-to-one (2:1) buffers or complex buffers (41, 44, 48):

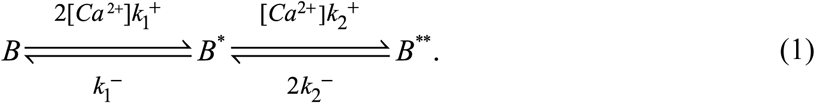

Here *B, B*^*^ and *B*^**^ denote respectively the free, partially bound, and fully Ca^2+^-bound states of the buffer, and 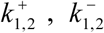 are the Ca^2+^-buffer binding/unbinding rates for each buffer state. Following previous modelling work (25), we will consider a semi-infinite domain bounded by a flat plane representing the cytoplasmic membrane, which contains one or more Ca^2+^ channels. Further, we assume zero flux boundary condition for Ca^2+^ and buffer on the flat plane, so the reflection symmetry allows to extend the domain to infinite space, while doubling the current strength, which places the Ca^2+^ current sources inside the domain (24, 25, 31). Denoting free Ca^2+^ concentration as *C*, and time differentiation as ∂_*t*_, we arrive at the following reaction-diffusion system for the concentrations of all reactants (48):

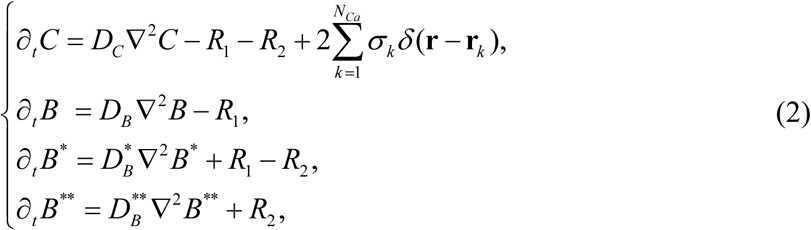

where the reaction terms are given by

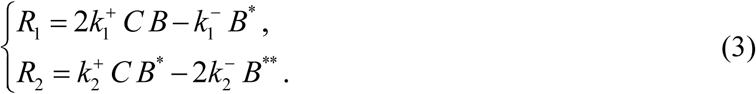

In Eq. 2, the point channel-source strengths are *σ*_*k*_ = *I*_*Ca,k*_/(2*F*), where *I*_Ca,k_ are the amplitudes of individual open Ca^2+^ channels located at positions **r**_*k*_, *F* is the Faraday constant, and *N*_Ca_ is the number of Ca^2+^ channels. As in the simple-buffer case (25, 26, 37, 38, 49, 50), there are two conservation laws for the total buffer and the total Ca^2+^ concentrations:

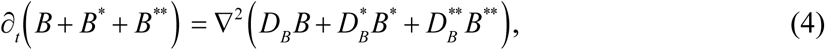

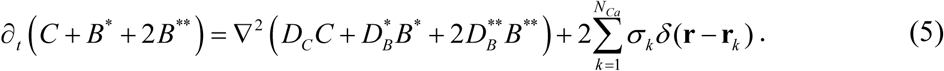

Since we are interested in equilibrium solutions, we obtain

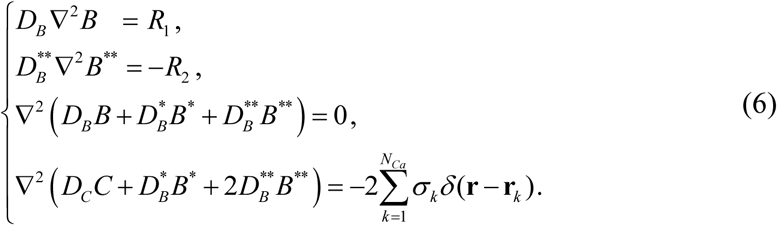

Here we assume that buffer diffusivity does not change when binding Ca^2+^ ions, 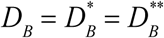 (this constraint is relaxed in the derivation of RBA in Supporting Material 1). In this case the two conservation laws in Eq. 6 have the following solution (25, 26, 31, 37, 48-50):

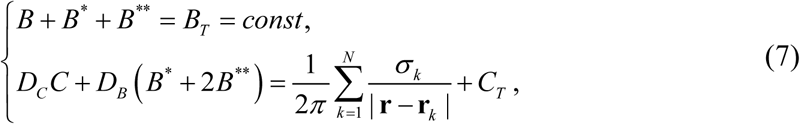

where constants *B*_T_ and *C*_T_ are related to the total (bound plus free) buffer and Ca^2+^ concentrations respectively, infinitely far from channel:

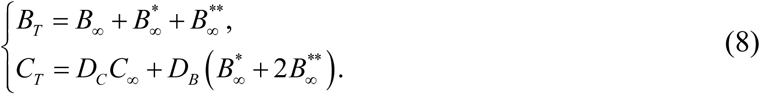

Here *X*_∞_ denote the concentrations of reactants *X* infinitely far from the channel, where reactions given by Eq. 3 are at equilibrium. Therefore, all background buffer state concentrations are uniquely determined by the background [Ca^2+^], *C*_∞_, through equilibrium relationships

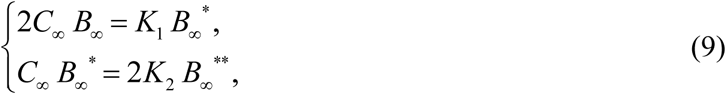

where *K*_1,2_ are the affinities of the two reactions in Eqs. 1, 3, given by 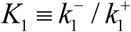 and 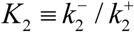.

We now restrict to the case of a single Ca^2+^ channel of source strength *σ* = *I*_*Ca*_/(2*F*) at the origin, and look for spherically symmetric solutions, which turns Eq. 6 into a system of ODEs, with the spherically symmetric Laplacian given in terms of the distance from the Ca^2+^ channel, 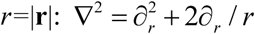:

We non-dimensionalize this problem using an approach analogous to the one we used for the simple buffer case in (37), which is a slightly modified version of the non-dimensionalization introduced by Smith et al. (25) and also used in (38, 48). Namely, we normalize Ca^2+^ and buffer concentrations by the affinity of the 2^nd^ binding step and the background buffer concentration, respectively:

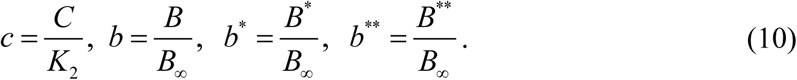

We also re-scale the spatial coordinate (*r*/*L* → *r*) using the scale parameter that depends on the strength of the Ca^2+^ current,

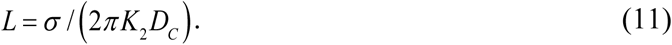

This transforms Eqs. 6,7 to the form

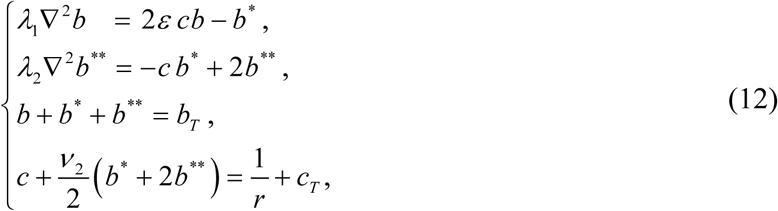

where *b*_T_ and *c*_T_ are the non-dimensional versions of the integration constants given by Eq. 8, related to the total buffer and [Ca^2+^] infinitely far from the channel (note that in our non-dimensionalization *b*_∞_=1):

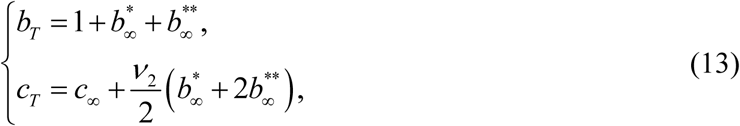

with dimensionless parameters

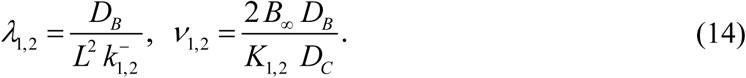

Along with *c*_∞_, parameters λ_1,2_ and *v*_1,2_ completely specify the model system. Here *λ*_1,2_ are the dimensionless mobilities of the two buffer states, which depend on buffering kinetics and Ca^2+^ current amplitude through the length scale *L* (Eq. 11). They quantify the ratio between the rate of diffusion and the rate of Ca^2+^ influx and binding. Parameters *v*_1,2_ quantify the overall buffering strength, and equal the product of the relative buffer mobility, *D*_B_ / *D*_C_, and the two quantities characterizing buffering capacity at rest, 2*B*_∞_ */K*_1,2_. For the sake of simplicity, we will also use the following cooperativity parameters, which characterize the difference between the affinities and kinetics of the buffer’s two distinct Ca^2+^-binding sites:

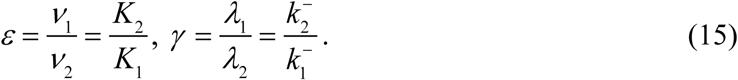

In the case of calretinin and calmodulin, the binding properties have been experimentally estimated (41, 45, 51), and the corresponding values of cooperativity parameters are given in Tables 4 and 5. These two Ca^2+^ buffer-sensors are characterized by highly cooperative Ca^2+^ binding, with *ε* <<1. In the results shown below, we will use the cooperativity parameters given by Eq. 15 to replace some of the four parameter given by Eq. 14. Namely, we will specify our model using either {*λ*_2_, *v*_2_, *ε, γ*} or {*λ*_1_, *λ*_2_, *ε, q*}, where *q*=1/(1+*v*_1_) is analogous to the parameter of the same name in the simple buffer case (37, 38).

**Table 1.** List of all new approximants, including the ansätze for *U* and *V* and the number of terms in the short-range and long-range solution expansions (Eqs. 19, 21) matched by each *ansatz*. Note that all *ansätze* automatically match the term of order O(*x*) in *U* (*U* ∼ 2*qx*) and the term of order O(*x*^2^) in *V* (*V* ∼ *q*^2^*x*^2^). The free parameter in the *U* ansatz (*α* or *A*) is found by matching terms of order O(*r*), while the free parameters in the *V* ansatz are found by matching terms indicated in the last column.

| Name | Ansatz in $U$ | Ansatz in $V$ | Params | $V$ accuracy |
| --- | --- | --- | --- | --- |
| PadéA | $U = \frac{2q}{A + r}$ | $V = \frac{q^2}{r^2 + b_1 r + b_2}$ | $N=3$<br>$(A, b_1, b_2)$ | $O(r), O(x^3)$ |
| PadéB | | | | $O(r^2), O(x^2)$ |
| ExpPadéA | $U = 2q \frac{1 - e^{-r/A}}{r}$ | | | $O(r), O(x^3)$ |
| ExpPadéB | | | | $O(r^2), O(x^2)$ |
| PadéExp | $U = \frac{2q}{A + r}$ | $V = q^2 \frac{1 - e^{-sr}(1 + sr)}{r^2}$ | $N=2$<br>$(A, s)$ | $O(r), O(x^2)$ |
| ExpExp | $U = 2q \frac{1 - e^{-r/A}}{r}$ | | | $O(r), O(x^2)$ |

**Table 2.**
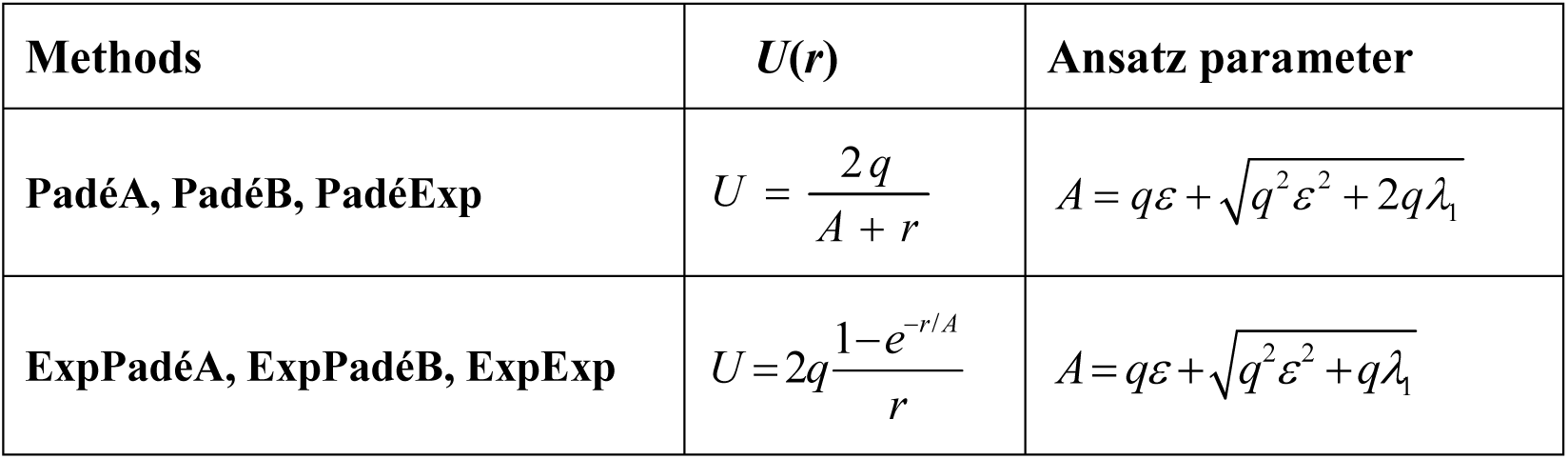
*Ansätze* parameters for the free buffer variable *U* as functions of model parameters *q, ε, λ*_1_, and *λ*_2_.

**Table 3.** *Ansätze* parameters for the fully bound buffer variable *V* shown in Table 1, as functions of model parameters *q, ε, λ*_1,2_, and parameter *A* from the corresponding *ansatz* for *U* shown in Table 2. For PadéB, ExpPadéB, PadéExp and ExpExp, the value of *b*_2_ or *s* is given by the real positive root of the cubic equation shown in the last column, whose closed-form solutions are given in Appendix A.

| Method | Equations for ansatz parameters (Eqs. 24, 25) |  |
| --- | --- | --- |
| PadéA, ExpPadéA | $b_1 = 2q[1 - q + \varepsilon(2q - 1)]$ | $b_2 = \frac{1}{4} \left( Aq + \sqrt{A^2 q^2 + 16b_1 A q \lambda} \right)$ |
| PadéB | $b_1 = \frac{b_2}{2\lambda_2} \left( \frac{2b_2}{Aq} - 1 \right)$ | $3\lambda_2 \left( \frac{b_1^2}{b_2} - 1 \right) + b_2^2 \frac{q-2}{qA^2} + \frac{b_1}{2} - b_2 = \frac{q(q-1)}{4}$ |
| ExpPadéB | | $3\lambda_2 \left( \frac{b_1^2}{b_2} - 1 \right) + b_2^2 \frac{2q-3}{2qA^2} + \frac{b_1}{2} - b_2 = \frac{q(q-1)}{4}$ |
| PadéExp, ExpExp | $4\lambda_2 s^3 + 3s^2 - 12/(qA) = 0$ | |

**Table 4.** Ca^2+^ binding properties of strongly cooperative buffers calretinin (CR) and calmodulin (CaM), as reported in (41, 51). Each CR molecule contains 5 binding sites, consisting of two identical cooperative pairs of Ca^2+^-binding sites and one independent non-cooperative site. CaM molecule consists of two independent domains (lobes), each binding two Ca^2+^ ions in a cooperative manner. Values of *λ*_2_ and *v*_2_ are calculated for Ca^2+^ current strength of *I*_Ca_=0.4 pA, total buffer concentrations of *B* =100 μM, buffer-Ca^2+^ mobility ratio of D_B_/D_Ca_=0.1, and D_Ca_=0.2μm^2^/ms.

| Parameter | $k_1^+$ | $k_2^+$ | $K_1$ | $K_2$ | $\varepsilon = \frac{K_2}{K_1}$ | $\gamma = \frac{k_2^-}{k_1^-}$ | $\lambda_2$ | $\nu_2$ |
| --- | --- | --- | --- | --- | --- | --- | --- | --- |
| Units | ($\mu\text{M ms}$)<sup>-1</sup> | ($\mu\text{M ms}$)<sup>-1</sup> | $\mu\text{M}$ | $\mu\text{M}$ | | | | |
| CaR (coop. 2×2) | 0.0018 | 0.31 | 28 | 0.068 | $2.4 \cdot 10^{-3}$ | 0.42 | $1.6 \cdot 10^{-3}$ | 294 |
| CaR (non-coop) | 0.0073 | -- | 36 | -- | -- | -- | -- |  |
| CaM (N-lobe) | 0.1 | 0.15 | 26.6 | 6.6 | 0.248 | 0.372 | 0.323 | 3.03 |
| CaM (C-lobe) | 0.004 | 0.01 | 10 | 0.93 | $9.3 \cdot 10^{-2}$ | 0.23 | 0.68 | 21.5 |

We now restrict our analysis to the case of zero background [Ca^2+^], relegating more general results to Supporting Material 1. With this simplification, *c*_∞_=*c*_T_=0 and *b*_∞_*=b*_T_*=*1, therefore Eq. 12 becomes

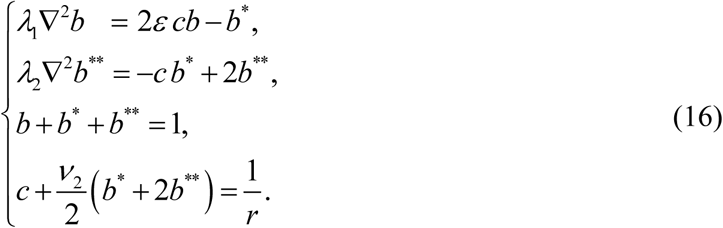

This system posed a challenge since it represents a non-linear and singular problem on an infinite domain. Further, most stationary approximations developed for the case of a simple 1:1 Ca^2+^ buffer cannot be extended to complex 2:1 Ca^2+^ buffers, with the exception of lowest-order RBA, which assumes that the reaction is at equilibrium in the entire domain (48). In Supporting Material 1 we derive RBA using the new non-dimensionalization presented above, slightly generalizing the expressions in (48). As is the case for a simple 1:1 Ca^2+^ buffer, RBA approximates the true solution very well within the parameter regime *λ*_1,2_<<1 (25). However, this fast buffering parameter regime has a complex interplay with the cooperativity condition *ε* <<1. In fact, the accuracy in buffer concentration estimation is significantly reduced with increasing Ca^2+^ binding cooperativity, corresponding to decreasing *ε*. Reducing the unbinding rate ratio *γ* along with *ε* partially rescues RBA accuracy (48). This high sensitivity of RBA accuracy to buffer parameters calls for the development of other approximants. In the results shown below, the accuracy of newly developed approximants will be compared and contrasted with that of the RBA.

Note that the stoichiometric factors of 2 appearing in Eqs. 12-14, 16 could in principle be absorbed into the definitions of the reaction rate parameters. However, we retain them, since as was pointed out in (48), this improves consistency with the non-dimensionalization for the simple 1:1 buffer case adopted in (25, 37, 38), allowing to recover the latter simpler model as *ε*→1 and *γ* →1.

In all results shown below, closed-form approximations to solutions of Eq. 12 or Eq. 16 are compared to the numerical solutions computed using the relaxation method and cross-validated using the shooting method; for the relaxation method we used CalC (Calcium Calculator) software, version 7.9.6 (http://www.calciumcalculator.org) (52).

## RESULTS

### Equilibrium Ca^2+^ nanodomain: power series interpolation method

We begin by presenting the power series interpolation method developed recently for the case of simple buffers with one-to-one Ca^2+^ binding stoichiometry (37, 38), which we will now generalize to the case of 2:1 Ca^2+^ buffers with two binding sites. This method involves finding simple *ansätze* that interpolate between the solution’s Taylor series in powers of distance from the channel location, *r*, and the asymptotic power series expansion of the solution in terms of the reciprocal distance from the channel location, *x*=1/*r*. We will refer to these two series as the short-range (low-*r*) and long-range (high-*r*) series.

We start with the non-dimensionalized form of the system for complex buffer, Eq. 16, and make a substitution *U* = (1− *b*) / *ε*, and *V* = *b*^**^ / *ε* to slightly simplify these equations. Eliminating the partially-bound buffer concentration variable using the buffer conservation law *b*^*^ = 1− *b* − *b*^**^ = *ε* (*U* −*V*), Eq. 16 is transformed to

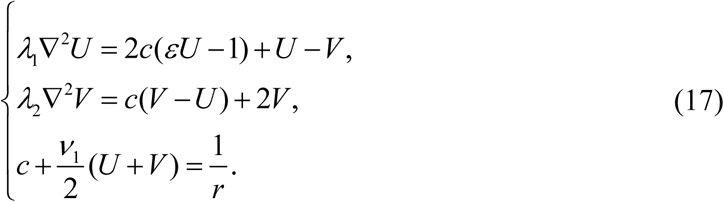

Next, we eliminate the Ca^2+^ concentration *c* using the Ca^2+^ conservation law in the last equation to obtain

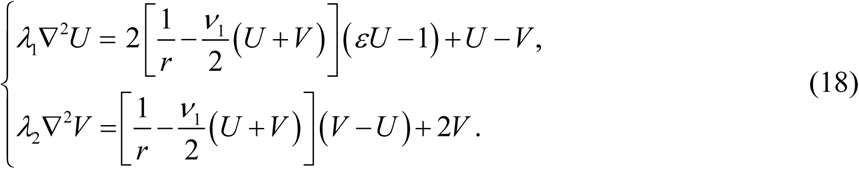

This system has only a regular singularity at *r*=0 and does have a solution analytic at *r*=0, representing the physical nanodomain solution that we are seeking. Using the Frobenius-like approach we find the following Taylor series expansions in *r* for both *U* and *V*:

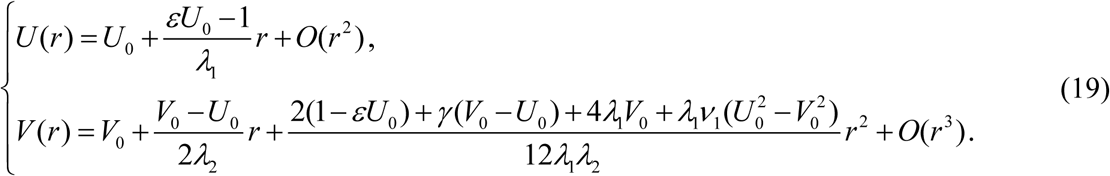

Parameters *U*_0_ and *V*_0_ are determined by the concentrations of free and fully bound buffer at the channel location, *r*=0; these values are finite and non-zero, but unknown *a priori*. Thus, *U*_0_ and *V*_0_ are important unknowns of the problem, to be determined by our approximation procedure. Because *U*_0_ and *V*_0_ are unknown, Eq. 19 only provides the relationships between the coefficients of these Taylor expansions, rather than coefficients themselves. For example, denoting the 1^st^-order Taylor coefficients in Eq. 19 as *U*_1_ and *V*_1_, we obtain the constraints *U*_1_=(*εU*_0_−1)/ *λ*_1_ and *V*_1_=(*V*_0_−*U*_0_)/(2*λ*_2_), which we will use to determine some of the free parameters of each approximant considered further below.

In order to obtain the long-range asymptotic series expansion of the solution, we make a coordinate mapping *x* ≡ 1/ *r*, transforming Eq. 18 to the form

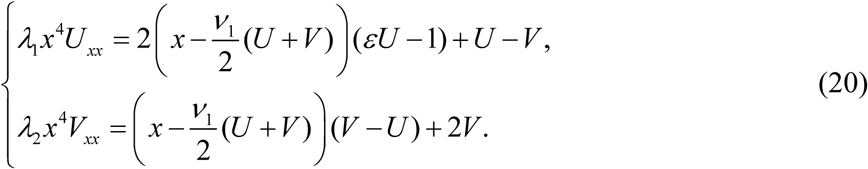

This system has a unique asymptotic power series expansion near *x=*0 satisfying boundary conditions at *x*→0^+^ (i.e. *r*→+∞), namely *U*(*x*=0^+^)=0, *V*(*x*=0^+^) =0. Up to terms of order *x*^3^, this asymptotic series expansion can be obtained by simply equating the right-hand sides of Eq. 20 to zero, which yields

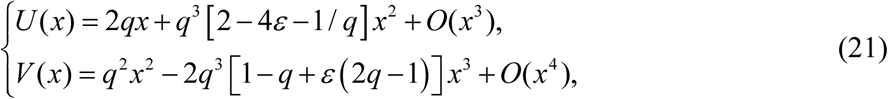

where we introduced the parameter *q* = 1/ (1+ *v*_1_) for the sake of simplicity. It is important to note that the leading term in the *V*(*x*) long-range expansion is of order O(*x*^2^), in contrast to *U*. This is intuitive, since *V* represents the double-bound buffer state, which decays faster than [Ca^2+^] or [B^*^] as [Ca^2+^]→0 with *x*→0^+^ (i.e. *r*→+ ∞). Note however that this is not the case when background [Ca^2+^] is not zero; this more general case is considered in Supporting Material 1. Parenthetically, we also note that the right-hand sides of Eq. 20 contain all reaction term, which RBA sets to zero. Therefore, given that the left-hand side of Eq. 20 is of asymptotic order O(*x*^4^), Eq. 21 must agree up to the given order O(*x*^3^) with RBA.

We will now consider simple *ansätze* whose series expansions simultaneously match leading terms of the low-*r* (short-range) series and the low-*x* (long-range, high-*r*) series given by Eqs. 19, 21. Inspired by the simple buffer case (37), we seek *ansätze* for *U* and *V* that combine Padé approximants (rational functions) and exponential functions. Below we list these *ansätze* for *U* and *V*, along with the corresponding short-range and long-range series representations. Our approximations are based on pair-wise combinations of these *U* and *V ansätze*, as summarized in Table 1. With a slight abuse of notation, we use the same function name (*U* or *V*) whether it is expressed as a function of distance *r*, or its reciprocal *x*:

- Padé *ansatz* for *U*, containing one free parameter, *A*:

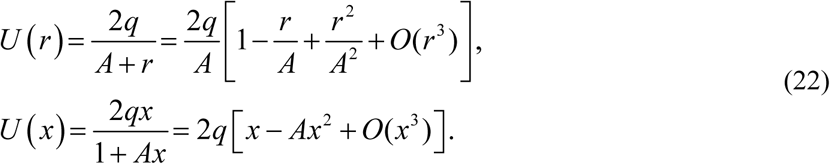
- Exponential *ansatz* for *U*, which also depends on one free parameter *A*:

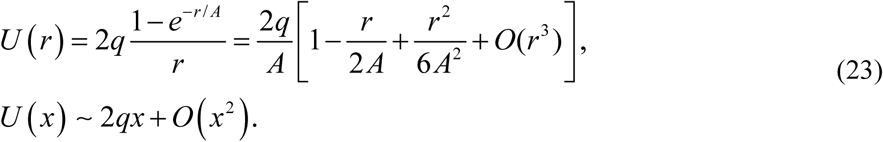
- Padé *ansatz* for *V*, which depends on two free parameters, *b*_1_ and *b*_2_:

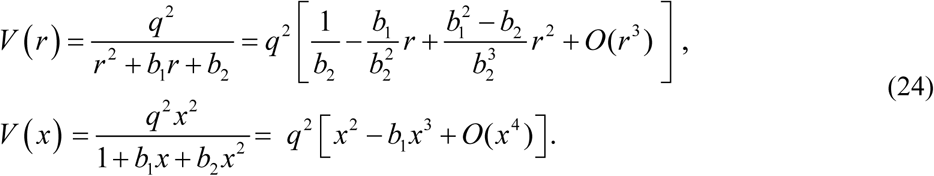
- Exponential *ansatz* for *V*, which depends on one free parameter, *s*:

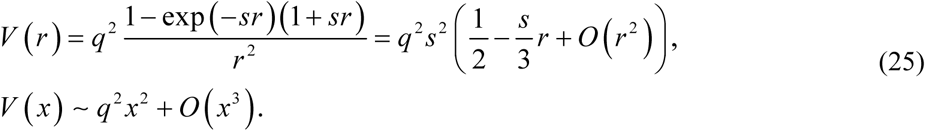

Note that all of these *ansätze* are analytic at *r*=0, and that in the limit *r*→+ ∞ (*x*=1/*r*→ 0^+^), they automatically match the leading non-zero term in the asymptotic series expansion of the solution given by Eq. 21: *U*(*x*)= 2*qx* + O(*x*^2^), *V*(*x*)=*q*^2^*x*^2^ + O(*x*^3^). Additionally, all *ansätze* satisfy appropriate physical constraints. Namely, imposing the condition *A* > 0 guarantees that *U* is positive and monotonically decreasing to 0 as *r*→ +∞ for each *ansatz*, and therefore *b*=1 at infinity, since *U* = (1− *b*) / *ε*. This agrees with the observation that the free buffer concentration is increasing monotonically from *b*_0_>0 at the channel mouth to *b*=1 infinitely far from the channel. Further, *V* is also always positive given positive parameters *b*_1_, *b*_2_, and *s*, and is monotonically decreasing to *V=*0 as *r*→ +∞, therefore *b*^*\*\**^=0 at infinity (recall that *V* = *b*^**^ / *ε*). This agrees with the fact that the fully bound buffer concentration is bounded and equals to zero infinitely far from the Ca^2+^ channel, where [Ca^2+^]=0.

We match the free parameters in the above approximants following the same interpolation method as in the case of a simple 1:1 Ca^2+^ buffer (37, 38). Namely, the unknowns are *U*_0_ and *V*_0_ in Eq. 19, plus either 2 or 3 parameters characterizing a particular approximant, as listed in Table 1. Therefore, 4 or 5 constraints are needed to find these unknowns. The first 4 constraints are obtained by matching the first two terms (of order O(1) and O(*r*)) in the short-range series for both *U* and *V*, as given by Eq. 19. For the 3-parameter approximants, the final 5^th^ constraint is needed, which comes from matching one additional term in the short- or the long-range series, as specified in the last two columns of Table 1. One obtains an algebraic system of 4 or 5 equations for the *ansatz* parameters, which are readily solvable in closed form.

Tables 2 and 3 show the exact expressions we obtain using this method for the free *ansatz* parameters in terms of the model parameters {*λ*_1_, *λ*_2_, *q, ε*}, except for *b*_2_ and *s* defined by solutions of cubic equations shown in the last column of Table 3, whose roots are given in closed form in Appendix A. Once *b*=1− *εU* and *b*^**^=*εV* are determined using these approximants, the partially bound buffer concentration *b*^*\**^ and Ca^2+^ concentration *c* can then be determined from *b* and *b*^*\*\**^ through the conservation laws in Eq. 16.

We will now illustrate this series interpolation method more concretely using the ExpPadéA approximant as an example. This *ansatz* is formed by combing Eq. 23 for *U* and Eq. 24 for *V*. Then, as indicated in Table 1, we constrain the values of *ansatz* parameters using terms of orders O(1) and O(*r*) in Eq. 19 for both *U* and *V*, and term of order O(*x*^3^) in Eq. 21 for *V* (recall once again that all *ansätze* automatically match the term of order O(*x*) in *U* and the term of order O(*x*^2^) in *V*). Therefore, we obtain 5 constraints for 5 unknowns (three parameters in ExpPadéA *ansatz*, plus *U*_0_ and *V*_0_):

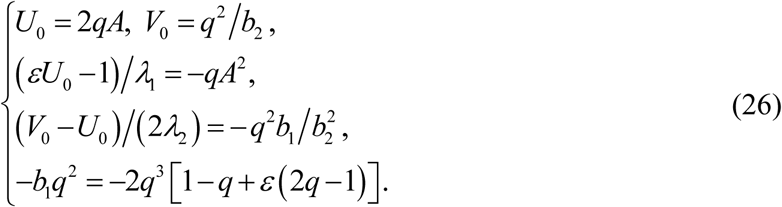

The solution of this system is given in Tables 2-3. Note that the 3^rd^ equation in this system leads to a quadratic equation for *b*_2_, whose solution is given in Table 3.

### Accuracy in approximating buffer and Ca^2+^ concentrations

As a crude demonstration of the performance of our new *ansätze*, Figure 1 shows our approximants for 4 select combinations of model parameters, with each column presenting the results for all concentration variables (*b, b*^*^, *b*^**^, and *c*), for a particular set of values of *λ*_2_, *v*_2_, *γ*, and *ε*, as labeled in the panel titles. The accurate numerical results are shown as thick grey curves. Since the expressions for the free buffer *b* (specified by *U*) are identical for PadéA, PadéB, and PadéExp approximants (see Table 2), the corresponding approximation is labeled as U-Padé, and shown as a single *dashed green curve* in the top panels of Fig. 1. Similarly, *b* approximants for ExpPadeA, ExpPadeB and ExpExp are also identical, and are labeled U-Exp and shown as a *dashed magenta curves* in the top panels. Only the best 5 approximants are shown for each parameter combination for variables *b*^*^, *b*^**^, and *c* in rows 2-4 of Figure 1 (out of a total number of 7 approximants combining the 6 *ansätze* in Table 1, plus RBA).

**FIGURE 1.**
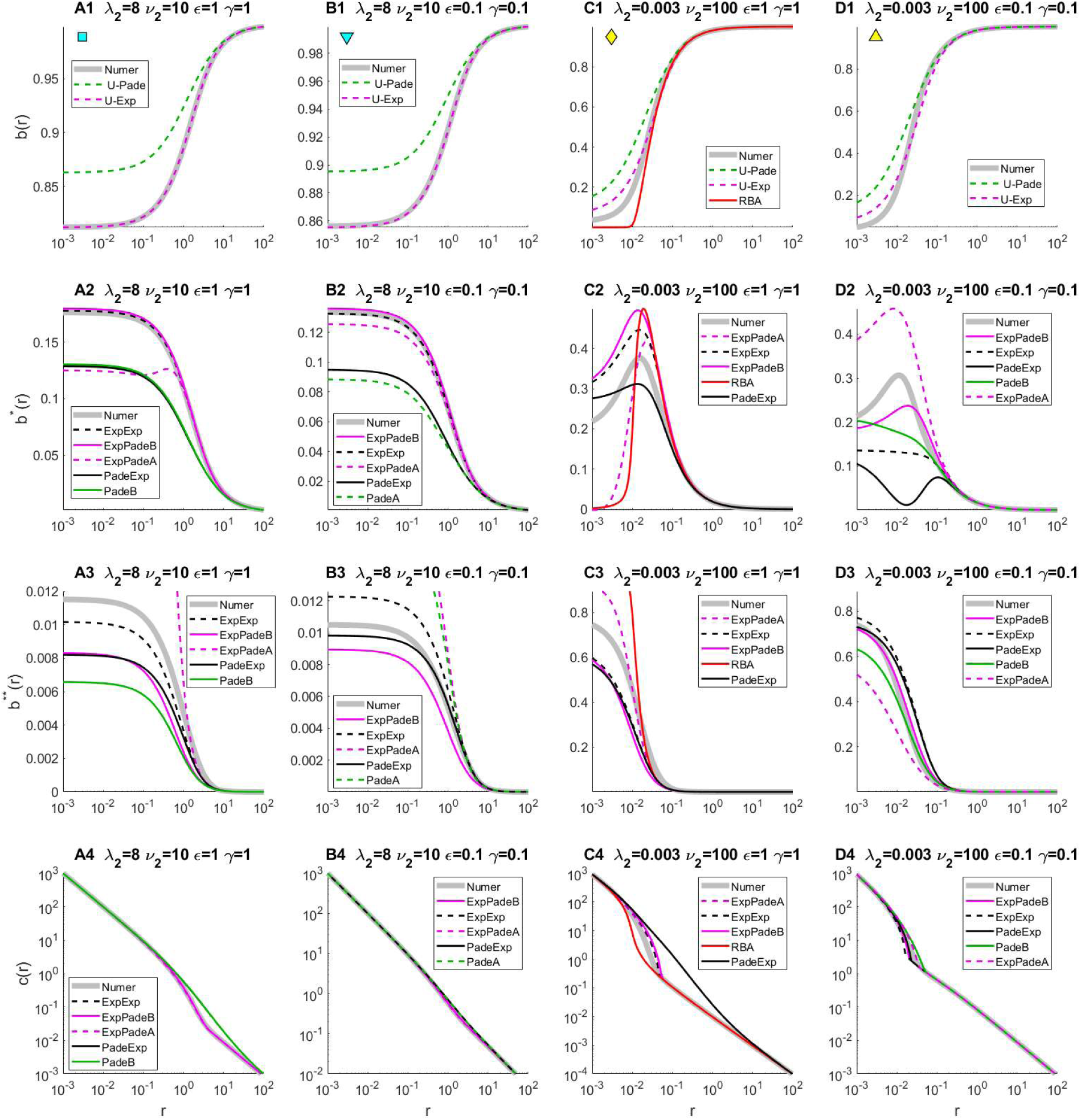
Approximations of equilibrium free buffer (top row), partially bound buffer (2^nd^ row), fully bound buffer (3^rd^ row) and Ca^2+^ concentrations (bottom row), as functions of distance from the Ca^2+^ channel, for 4 distinct choices of model parameters *λ*_2_, *v*_2_, *γ*, and *ε* indicated in the panel titles. The distinct curves mark the series interpolants shown in Table 1: PadéA (*green curve*s), PadéB (*dashed green*), ExpPadéA (*magenta*), ExpPadéB (*dashed magenta*), PadéExp (*black*), ExpExp (*dashed black*), and RBA (*red*). Since these approximants involve only 2 distinct *ansätze* for the free buffer variable *U* (see Tables 1,2), the latter are labeled as U-Pade (*dashed green*) and U-Exp (*dashed magenta*) in panels A1, B1, C1, D1. *Grey curves* show the accurate numerical simulations. A subset of 5 best methods is shown for each parameter combination. The accuracy of some approximants is sufficiently high for the curves to completely overlap with the numerical solution on the given scale, and hence the difference between the curves is hard to resolve by eye.

As will be elucidated further below (see Figs. 3-6), the parameter regimes we selected in Fig. 1 are not optimal for the *ansätze* we introduce. Nevertheless, even for the chosen sub-optimal parameter combinations, a decent qualitative agreement with the accurate numerical solution is achieved by at least one of the *ansätze*, with higher accuracy achieved for the first two parameter combinations in Fig. 1A1-4, B1-4. We observe that RBA can compete with the newly presented approximants only when diffusivity *λ*_2_ is very small (Fig. 1C1-C4); therefore, RBA is not shown for the other three parameter choices. Note the difference in scales in the different panels of Fig. 1: some of the apparent large discrepancies for *b*^*^ and *b*^**^ involve relatively small absolute differences. The accuracy of several of the newly presented approximants is sufficiently high for the curves to completely overlap with the numerical simulations. Therefore, the series interpolation method achieves significant improvement of approximation accuracy for a wide range of model parameters, as compared to RBA.

**FIGURE 2.**
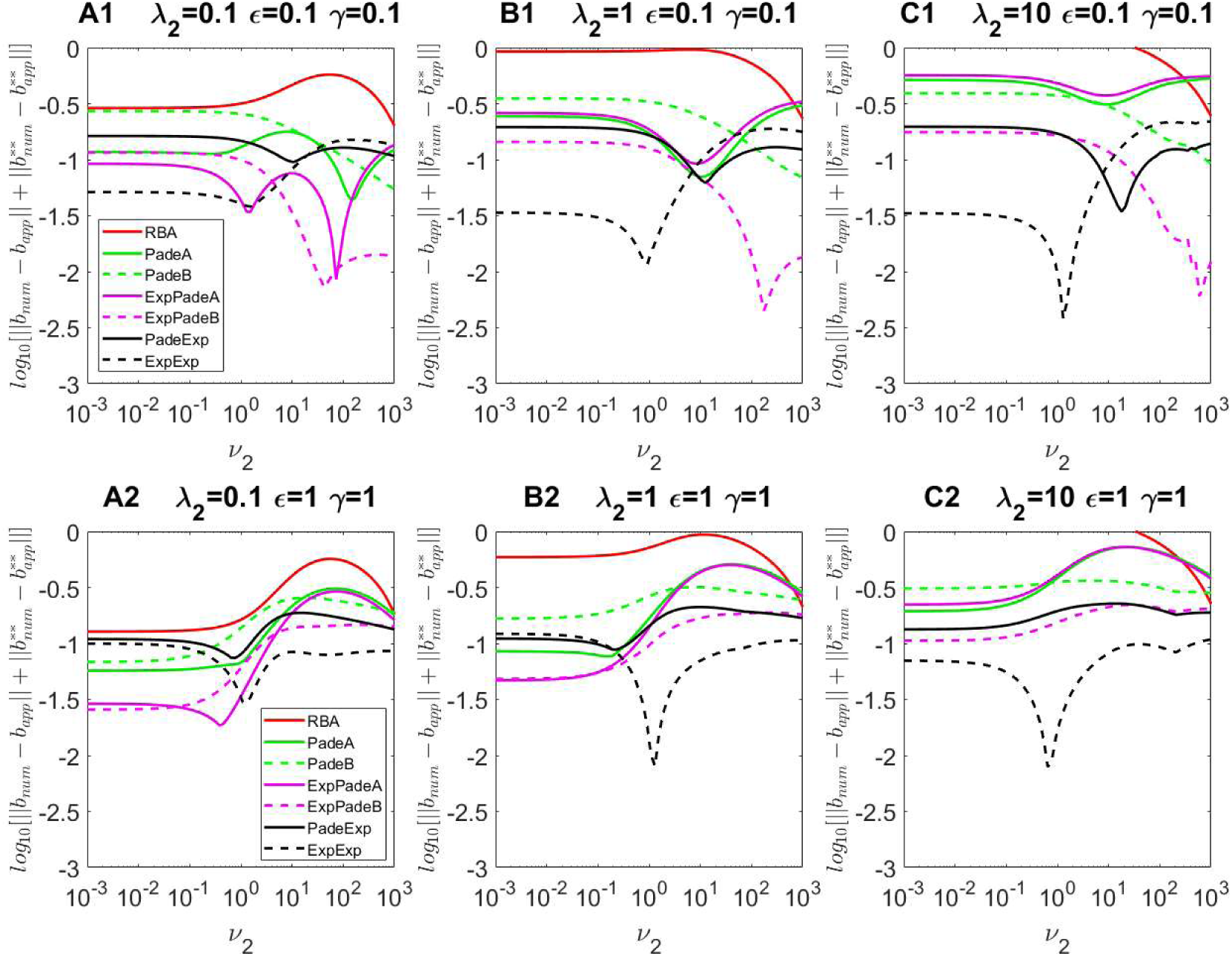
Accuracy comparison of the series interpolant approximations for the equilibrium free and fully bound buffer concentrations: PadéA (*green curves*), PadéB (*dashed green*), ExpPadéA (*magenta*), Exp-PadéB (*dashed magenta*), PadéExp (*black*), and ExpExp (*dashed black*). RBA error is also plotted for comparison (*red curves*). All panels show the log_10_ of the sum of average errors of approximating concentrations of free buffer (*b*) and fully bound buffer (*b*^**^) computed using Eq. 27, as a function of buffering strength *v*_2_ ranging from 10^−3^ to 10^3^, for 3 distinct choices of parameters *λ*_2_, *ε* and *γ*, as indicated in panel labels.

**FIGURE 3.**
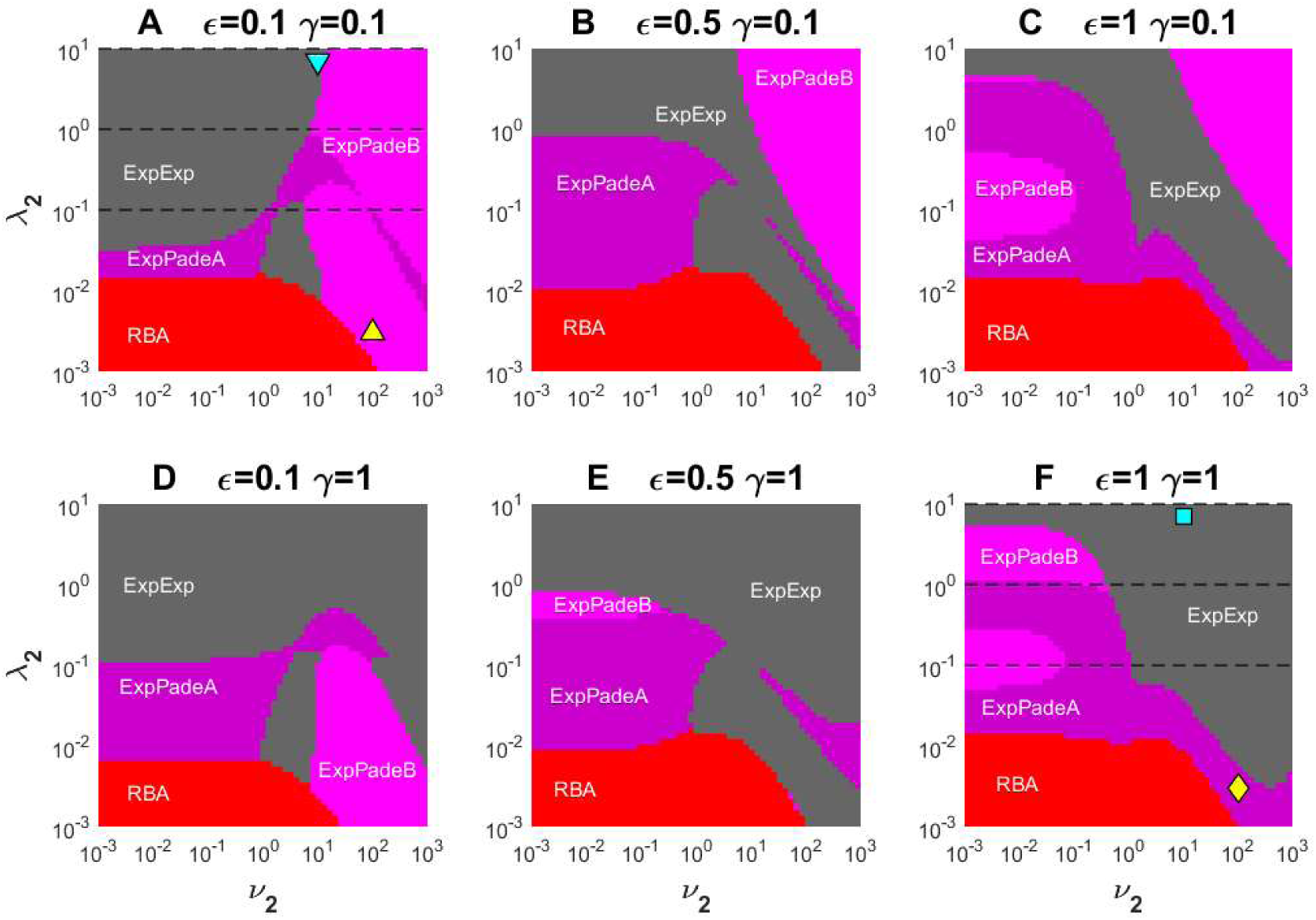
Comparison of parameter regions where a given approximant outperforms the rest in estimating the combined errors of free and fully bound buffer concentration approximations in the (*v*_2_, *λ*_2_) parameter plane, according to the error measure given by Eq. 27, with 6 different choices of cooperativity parameters *ε* and *γ*: (A) *ε*= *γ* =0.1; (B) *ε* =0.5, *γ* =0.1; (C) *ε* =1, *γ* =0.1; (D) *ε* =0.1, *γ* =1; (E) *ε* =0.5, *γ* =1; (F) *ε*= *γ*=1. Each color in A through F marks the parameter region of best performance for the following approximants: RBA (*red*), ExpPadéA (*dark magenta*), ExpPadéB (*light magenta*), and ExpExp (*gray*). Yellow and cyan symbols mark parameter point corresponding to simulations in Figure 1, where the free and fully bound buffer concentrations are plotted separately. Dashed lines mark the locations of parameter scans in Figure 2.

It is interesting to note that the partially-bound buffer concentration *b*^*^ is not necessarily monotonic with respect to distance from the origin, unlike the free and fully-bound concentration variables. Despite the simple functional forms of our *ansätze*, they do in fact reproduce this non-monotonic behavior: see for instance the ExpPadéB approximant in Fig. 1C2,D2.

Figure 1 shows that ExpPadéA, ExpPadéB, and ExpExp give more consistently accurate results, at least for the examined parameter sets. As in the simple buffer case, buffer approximations have the lowest accuracy near the channel, and the greatest accuracy far from the channel, since buffer concentrations at the channel location are unknown, while the long-range asymptotic behavior of the true solution is known, and given by Eq. 21. In contrast, the differences between distinct Ca^2+^ approximations and the numerical solution are shown on a logarithmic scale, and are more pronounced at intermediate distances from the channel, due to the dominance of the free source term 1/*r* near the channel, which is the same regardless of model parameters (Eq. 17). Note also that the [Ca^2+^] traces shown in the last row of Figure 1 are obtained using the Ca^2+^ conservation law (Eq. 17), based on inexact approximations for *b* and *b*^**^. Therefore, no direct physical constraints on Ca^2+^ are imposed by this procedure. For specific parameters regimes, this may result in negative values of [Ca^2+^] sufficiently far from the channel, where the corresponding true concentration values are positive but small. This indeed happens for very large values of buffering strength, *v*_1,2_ ≥ 100. When this occurs, we use the RBA approximation (see Supporting Material 1) as a lower bound on Ca^2+^, since RBA becomes accurate sufficiently far from the channel for any model parameter values, as we noted above. Moreover, our extensive numerical investigation leads us to conjecture that RBA in fact represents a sub-solution (a pointwise lower bound) for the true [Ca^2+^]. This imposed truncation of [Ca^2+^] from below using RBA helps us correct the errors in estimating Ca^2+^ at larger distances when buffering is very strong (see for instance Fig. 1C4). Even in cases where negative [Ca^2+^] values are replaced with RBA values, the accuracy of the new methods at closer distances are significantly improved compared to the RBA solution, as is the case for instance for the parameters in Fig. 1C4.

Examining approximation behavior for several example parameters combinations is insufficient to unveil the complicated parameter-dependent accuracy of these approximations. Therefore, following prior work (25, 37, 38, 48), we will now systematically explore the parameter-dependence of the absolute deviation between the given approximation and the accurate numerical solution, using the following norm, similar but slightly different from the norm used in the case of simple buffer (37, 38):

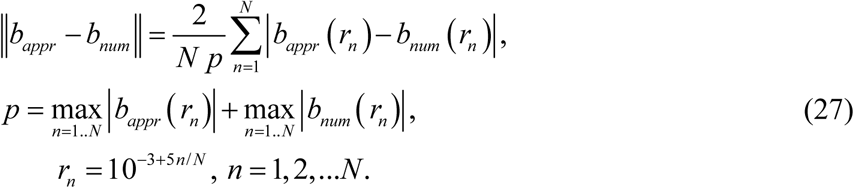

The deviations are computed on a set of *N* =100 points spanning 5 orders of magnitude of distance *r*, from 10^−3^ to 10^2^, on a logarithmic scale. Therefore, apart from the normalization factor *p* in the denominator (explained below), these are effectively L^1^ norms weighted by 1/*r*, which requires a lower distance cut-off, set to *r*_min_=10^−3^. The heavier weighting of short distances is justified by two reasons: (1) as we mentioned, our method has the greatest error at the channel location, and (2) the accuracy close to the channel is more important for actual biophysical modeling applications. We use the same error measure for approximating bound buffer state *b*^**^ as for *b*. Given the difference in absolute magnitude of *b*^**^ and *b*, we normalize by the maximal concentration *p* in the denominator of Eq. 27 to make it an even more stringent accuracy measure: as Fig. 1 illustrates, *b*^**^ can be quite small in certain parameter regimes, as compared to the free buffer *b*, which always approaches 1 as *r*→ +∞.

Since *b*^*^ and *c* are uniquely determined by *b* and *b*^**^ through the conservation law (Eq. 16), in all figures below we will examine the sum of errors for *b* and *b*^**^, instead of analyzing the different concentration fields individually. In Figure 2 we show a systematic comparison of the accuracy of the new approximants by plotting such sum of errors in *b* and *b*^**^ for each approximant as a function of the buffering strength parameter *v*_2_ varying from 10^−3^ to 10^3^, for three different fixed values of the buffer diffusivity parameter *λ*_2_ (*λ*_2_=0.1, *λ*_2_=1, or *λ*_2_=10) and two combinations of cooperativity parameters (*ε, γ*). To reveal the impact of Ca^2+^-binding cooperativity on approximant performance, one choice of (*ε, γ*) values corresponds to a non-cooperative buffer (*ε*= *γ=*1, bottom panels in Fig. 2), while the other choice corresponds to a very cooperative buffer (*ε*= *γ=*0.1, top panels in Fig. 2). Combining the error measures of *b* and *b*^**^ allows us to infer a single best approximation for each given parameter combination. The error of RBA (*red curves*) is also included for the sake of comparison.

For most combinations of parameters examined in Figure 2, ExpPadéA, ExpPadéB, and ExpExp achieve the best accuracy compared to other approximants, which is consistent with the results show in Figure 1. For the non-cooperative case *ε*= *γ*=1 (bottom row of panels in Fig. 2), for sufficiently large values of *v*_2_ and *γ*_2_ the best approximating method is always ExpExp, and the average relative error is always below 10%, which is very good for such a simple approximation and such a stringent error measure. For the cooperative buffer case, *ε*= *γ*=0.1 (top row in Fig. 2), the individual error curves are getting more tangled, and the choice of best method is somewhat more complicated, but in general ExpExp achieves superior accuracy at smaller values of buffering strength *v*_2_. At larger values of *v*_2_, ExpPadéB becomes the best approximation method. Although RBA performs worse compared to other approximants for parameter conditions examined in Figure 2, the advantage of RBA for smaller values of *λ*_1,2_ will be revealed in the results presented next.

Figure 3 summarizes and extends the results presented in Figure 2, labeling the best approximants for a wide range of buffer mobility *λ*_2_ varying over 4 orders of magnitude, and buffering strength *v*_2_ varying over 6 orders of magnitude, for 6 fixed sets of cooperativity parameters *ε* and *γ* corresponding to each of the 6 panels. The selection of best approximant in Figure 3 is based on the minimal sum of errors of *b* and *b*^*\*\**^ approximations; the corresponding smallest error value is shown in Figure 4. As noted above, using this combined error measure helps in determining the single best approximation method for a given set of model parameters, recalling that *b*^*\**^ and *c* are uniquely determined by *b* and *b*^*\*\**^ (Eq. 16). Note that we exclude PádeExp, PádeA and PádeB methods in this comprehensive comparison: even though there are parameter regions where these 3 methods outperform others, these parameter regions are relatively small, and the accuracy advantage is not very significant. Figure 3 shows that there is still a significant portion of parameter space where RBA outperforms our newly developed methods, but as expected, this only happens for sufficiently small values of *λ*_1,2_. As Figure 4 shows, a qualitative accuracy within 10% is always guaranteed for all examined parameter combinations, and for some narrow parameter regimes the accuracy can be extremely high, with error reaching down to 0.025%.

**FIGURE 4.**
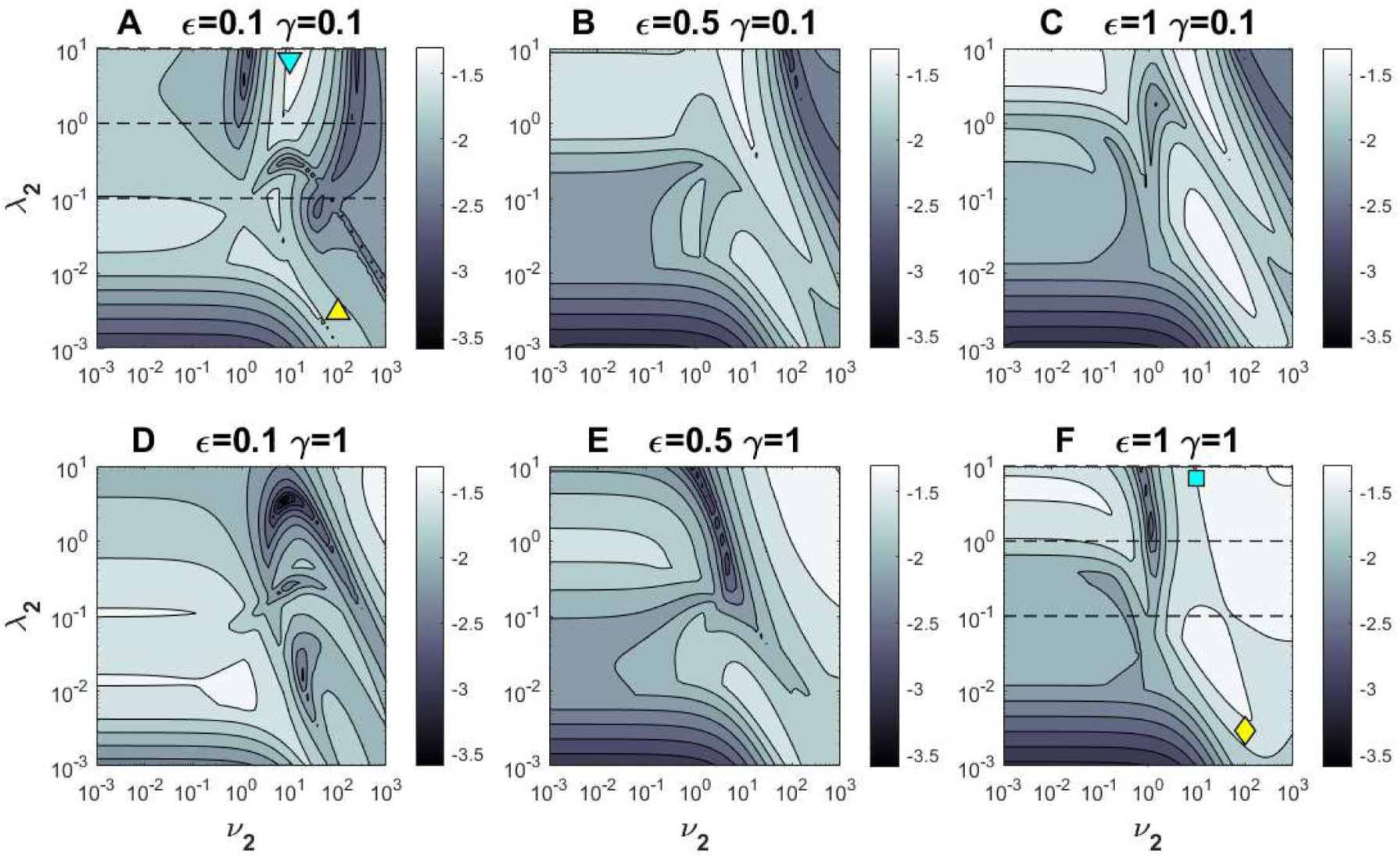
The smallest error in estimating the free and fully bound buffer concentrations in the (*v*_2_, *λ*_2_) parameter plane, according to the error measures given by Eq. 27, with *ε* and *γ* fixed to 6 different choices, as in Figure 3. The gray-scales in A through F indicate the log_10_ of the sum of average errors of the free and the fully bound buffer approximations (Eq. 27). Darker shades represent better accuracy, according to the error bars to the right of each panel.

Even though Ca^2+^ is uniquely determined from the buffer concentrations by the Ca^2+^ conservation law, it is still useful to look at the performance of different approximants in estimating [Ca^2+^] separately, since the latter is of obvious physical importance and has a different behavior as a function of distance from the channel. As noted above, close to the channel location [Ca^2+^] is dominated by the source term, 1/*r*, therefore we will modify the buffer error norm, Eq. 27, by taking the logarithm of [Ca^2+^] (37, 38, 48):

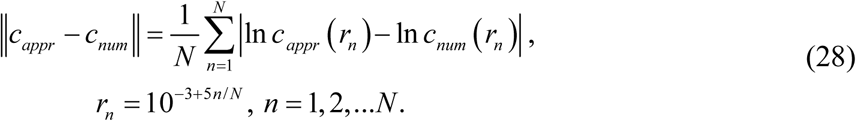

We note that qualitatively this norm has the same behavior as the relative norm used in (25). Figure 5 labels the approximants which minimize this error in estimating [Ca^2+^], with the corresponding error value shown in Fig. 6, using the same parameter combinations as in Figs. 3 and 4. Figure 5 shows that for any particular set of model parameters, the optimal approximants for [Ca^2+^] can be different from the optimal buffer approximant shown in Fig. 3, despite the fact that [Ca^2+^] is directly calculated from buffer concentrations. As discussed above, [Ca^2+^] approximant performance is more sensitive to its accuracy at intermediate distances, in contrast to the buffer error measure, which is the greatest in the channel vicinity. Therefore, the error in Ca^2+^ estimation measures the accuracy of our approximants at intermediate distance from the channel, while the error in buffer estimation reveals the method accuracy proximal to the channel location. This fact can also be observed in Figure 1.

**FIGURE 5.**
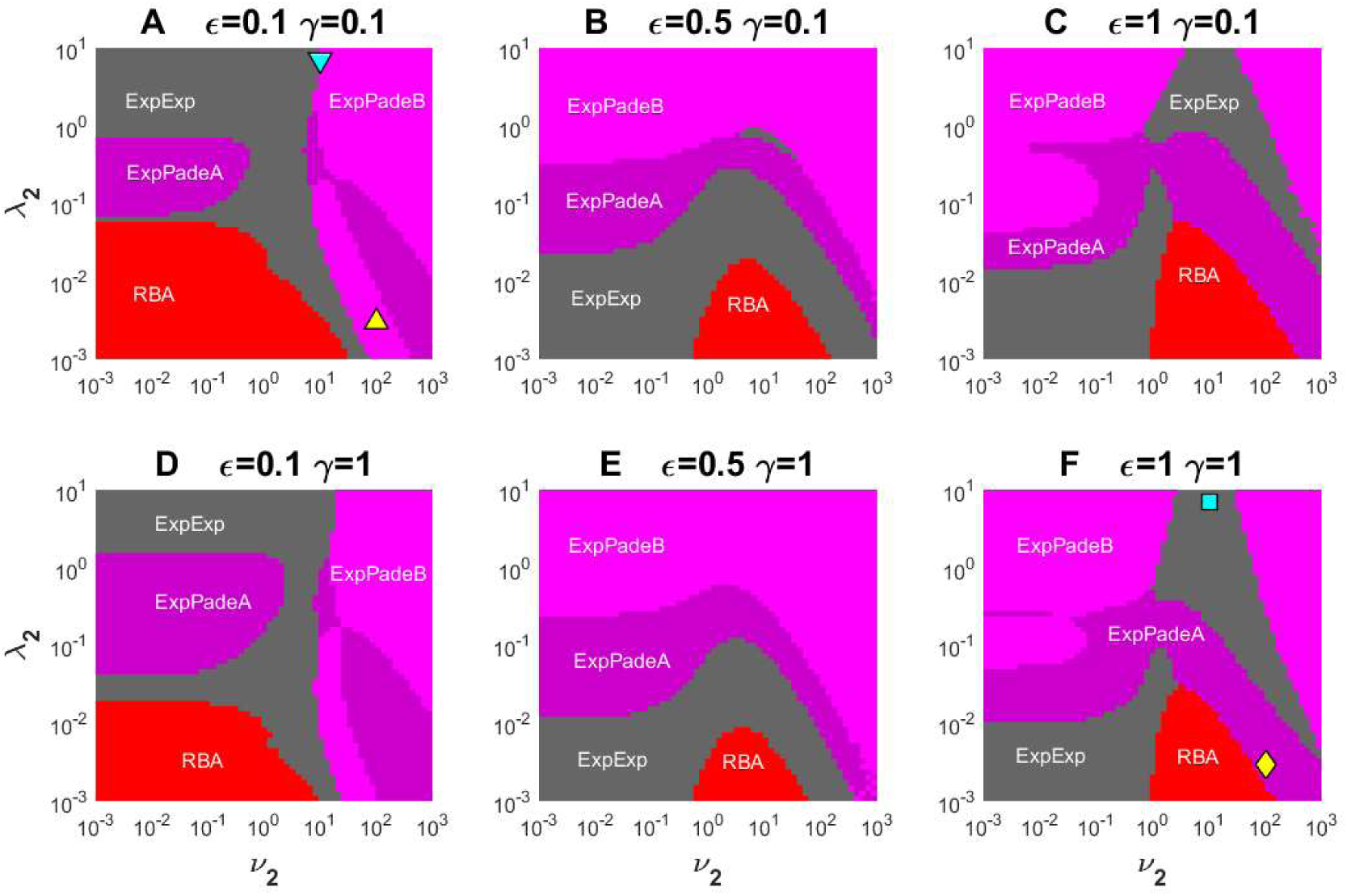
Comparison of parameter regions where a given approximant outperforms the rest in estimating [Ca^2+^] in the (*v*_2_, *λ*_2_) parameter plane, according to the error measure given by Eq. 28, with *ε* and *γ* fixed to 6 different choices, as in Figure 3. Each color in A through F marks the parameter region of best performance for the following approximants: RBA (*red*), ExpPadéA (*dark magenta*), ExpPadéB (*light magenta*), PadéExp (*black*), and ExpExp (*gray*). Yellow and cyan symbols mark parameter points corresponding to simulations in Figure 1.

**FIGURE 6.**
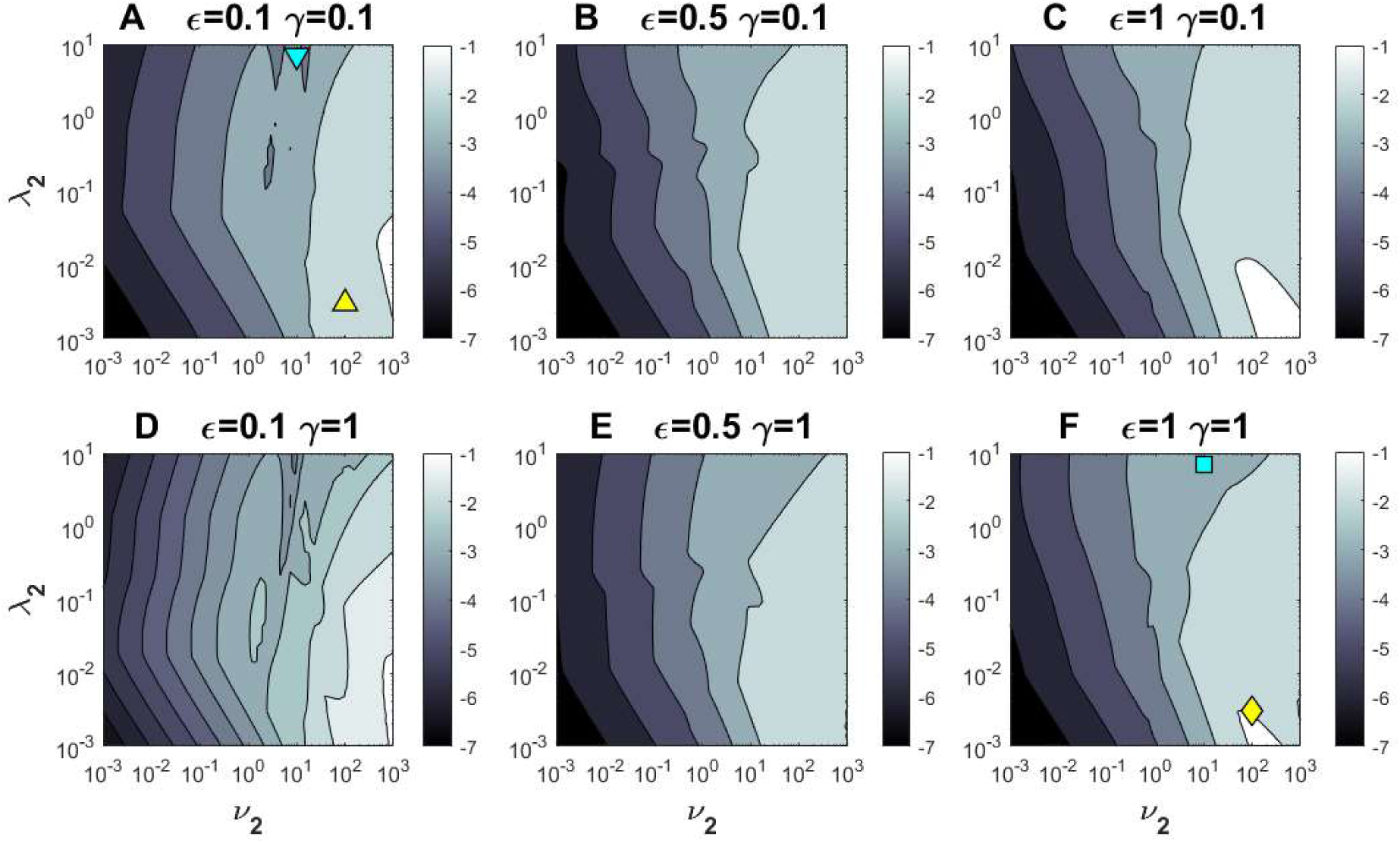
The error in estimating [Ca^2+^] in the (*v*_2_, *λ*_2_) parameter plane, obtained using the best approximant shown in Figure 5 for each parameter point, with *ε* and *γ* fixed to 6 different choices, as in Figure 5. All parameter choices and panel layout are identical with Figures 3-5. The grayscale in all panels indicates the log_10_ of error value given by Eq. 28, as indicated in scale bars to the right of each panel. Darker shade represents better accuracy.

Although PadéExp approximant is excluded from the comparisons shown in Figs. 5 and 6, it outperforms other methods in limited regions of parameter space corresponding to small *λ*_2_ and either very large or very small *v*_2_; however, even in those narrow parameter regions, the advantage of PadéExp is not very significant. Finally, we note that the uneven boundaries between accuracy levels in Fig. 6 do not indicate a numerical artifact, but are a consequence of the complicated shape of the boundaries of optimal performance regions that are shown in Fig. 5.

Finally, in order to evaluate whether our newly developed approximants achieve sufficient accuracy for parameters corresponding to real biological buffers, in Figure 7 we simulate the Ca^2+^ nanodomains in the presence of 100μM of Ca^2+^ buffer with the properties of either calretinin or one of the two lobes of calmodulin, shown in Table 4. For calretinin, we use parameter values reported by Faas et al. (41), while for calmodulin, we use reaction parameters that were carefully compiled from multiple biochemical studies by Ordyan et al. (51). As Figure 7 reveals, our newly developed method, ExpPadéA and ExpPadéB, work remarkably well for N-lobe or C-lobe of calmodulin: the curves for *b, b*^*\*\**^, and *c* corresponding to the approximations and the numerical simulations completely overlap at the chosen ordinate scale. For calretinin, ExpPadéB works the best, and demonstrates very reasonable accuracy. Although ExpPadéB fails to accurately describe the behavior of the single-bound calretinin concentration, it does accurately capture the order of magnitude of this concentration variable; further, note that the latter is much smaller than the free and fully bound buffer concentrations.

**FIGURE 7.**
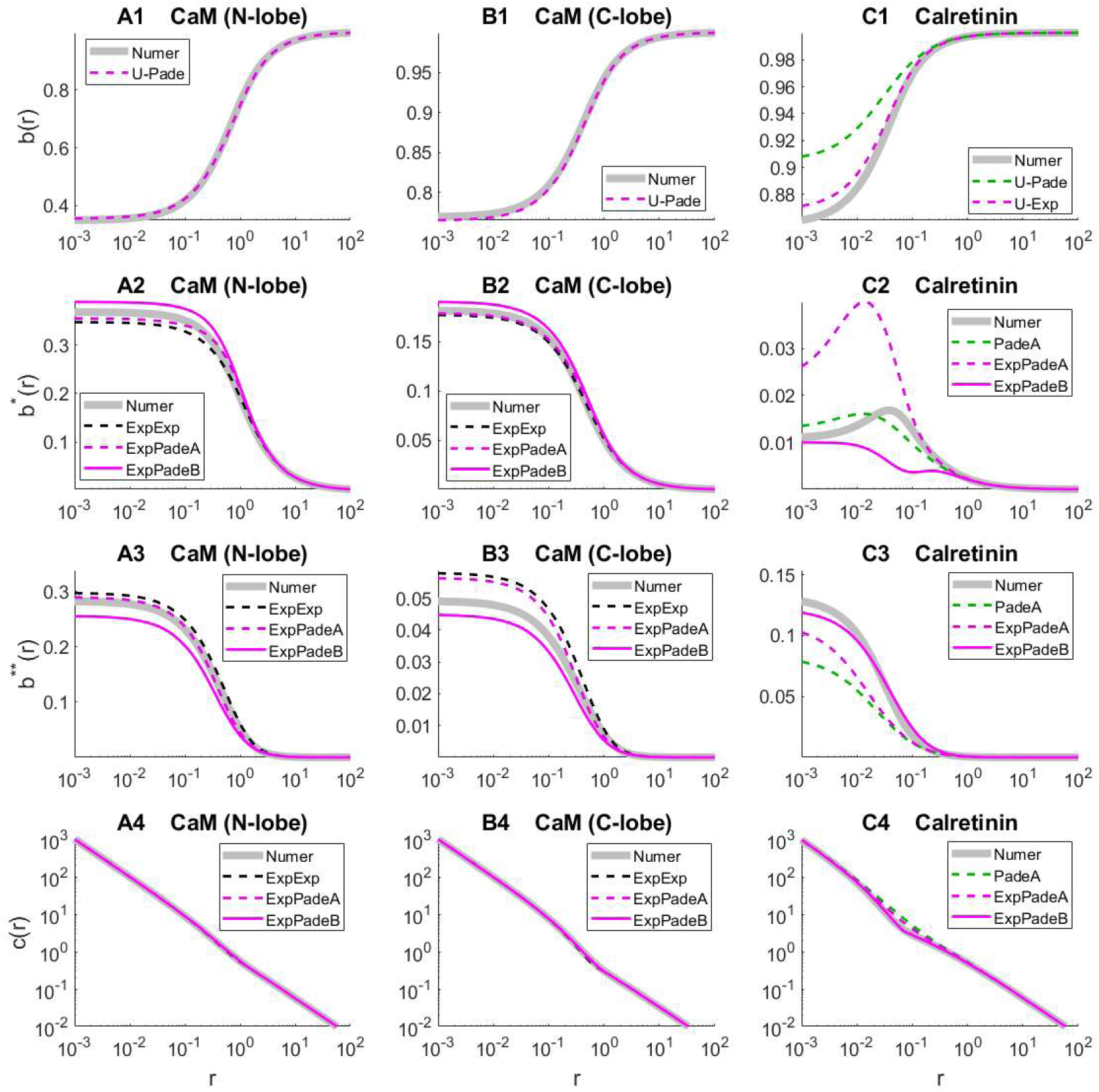
Approximation performance for the case of biological buffers, calmodulin N-lobe (A1-A4), calmodulin C-lobe (B1-B4), and calretinin (C1-C4), with parameters as in Table 4, corresponding to the current of *I*_Ca_=0.4 pA, and total buffer concentration of 100 μM. As in Figure 1, approximants of free buffer concentrations in panels (A1, B1, C1) are labeled as U-Exp and U-Padé (see Tables 1-2), while only the best approximations are shown for the other concentration variables: ExpPadéA (*solid magenta curve*), ExpPadéB (*dashed magenta*), PadéA (*dashed green*), and ExpExp (*dashed black*). Accurate numerical results are shown as thick gray curves.

Although the approximants we present do not allow to model the simultaneous impact of both lobes of calmodulin, since this would require generalizing our approach to buffer with 4 binding sites, the results obtained for the N-lobe alone are of value, since the N-lobe has much faster kinetics, and would reach a quasi-equilibrium state on short time scales compared to the much slower C lobe.

## DISCUSSION

We demonstrated that the series interpolation approach, first introduced for the case of 1:1 Ca^2+^ buffers in (38) and generalized in (37), can be extended to buffers with 2:1 Ca^2+^-binding stoichiometry, and we introduced several simple interpolants that combine rational and exponential functions. As summarized in Figures 3-6, and Figure S1 of the Supporting Material 1, these new approximants achieve reasonable accuracy in estimating equilibrium buffer and Ca^2+^ concentrations near an open Ca^2+^ channel in a wide range of relevant model parameters. Nevertheless, RBA, the only previously developed method for 2:1 buffers, is still superior for certain extreme parameter conditions corresponding to very small values of non-dimensional mobility parameters *λ*_1,2_. Compared with RBA, the new approximation methods show more uniform error dependence for several orders of magnitude of the relevant dimensionless parameters *λ*_2_, *v*_2_, *γ*, and *ε*. As Figure 4 shows, with these new approximants the average combined error for the free and fully bound buffer concentrations is within 10% for all examined parameter combinations, and Fig. 6 demonstrates similar maximal error in estimating Ca^2+^, albeit requiring truncation to ensure the physical constraints [Ca^2+^]>0 for large values of buffering strength. Figs. 3 and 5 illustrate that this accuracy level in the entire parameter range we considered is achieved with only 4 out of the total of 7 approximants, namely ExpPadéA, ExpPadéB, ExpExp, and RBA. Fig. 7 further shows that good qualitative agreement can be achieved even with more extreme model parameter values corresponding to calretinin or one of the two lobes of calmodulin, which correspond to parameter combinations shown in Tables 4 and 5.

Several functional forms other than the ones shown in Eqs. 22-25 were considered, but are not presented here due to either insufficient performance or lack of closed-form solutions for parameters. However, given the simplicity of the interpolating approximants we presented, improved *ansätze* could potentially still be found. This is particularly likely for the case of non-zero background Ca^2+^ concentration examined in the Supporting Material 1: only the simplest lowest-order interpolating approximants were consider in the latter general case.

Figs. 2-6 show that the accuracy profile of the approximants we introduced is highly non-trivial, with the error measure exhibiting large dips for certain parameter combinations. This is of potential interest and may reveal interesting properties of the underlying true solutions, deserving a careful investigation in the future.

Of course, practical use of the proposed approximants requires an algorithm for the choice of a particular *ansatz*, given a particular set of model parameters, without knowing the exact solution. Figs. 3,5 and Fig. S1 provide the basis for developing such an algorithm. Although the boundaries between parameter regions of best performance look complicated, a smaller subset of only three methods can allow one to develop a simple approximant selection algorithm, without sacrificing too much accuracy, as in the case of a 1:1 buffer (37). Here we note that, just like in the latter case, the relative accuracy comparisons of distinct algorithms summarized in Figs. 3-6, S1 depends on the particular norm that we have chosen for comparison to the true solution, given by Eqs. 27, 28.

There are many other directions for possible extensions and improvements of this work. For example, our approximants are only applicable to a single channel and a single buffer with two binding sites, whereas RBA allows an extension to an arbitrary number of channels and buffers (although the latter requires considerable increase in complexity). We note however that the methods we presented could be extended to the case of two distinct Ca^2+^ buffers with a single binding site each. We should also mention that we did not consider any Ca^2+^ sinks and the effect of finite channel pore radius. Including a linear homogeneous Ca^2+^ uptake mechanisms, along the lines of (49, 50), would greatly improve the utility of the developed approximations. Further, the utility of our approximants would be improved if one could find a method of estimating the method accuracy with respect to the chosen norms, without knowing the accurate numerical solution. For instance, one could examine whether barrier functions (sub- and super-solutions) could be used to establish the bounds on the approximant accuracy (54). Finally, the study of equilibrium concentration nanodomains assumes that the steady-state is established almost instantly and ignore the transient dynamics before the equilibrium is reached. However, this is not always the case (12). Therefore, the characteristic time needed to reach the steady state should be properly examined for a wide range of parameter values. Some related work on the time scale of the transients in reaction-diffusion systems can be found in (55). More generally, the fundamental mathematical analysis for the complex buffer case, along the lines of analysis in Appendix D of (37), is yet to be performed. For instance, we did not provide a rigorous proof that RBA is a sub-solution for [Ca^2+^] in this problem. Finally, we only considered the series interpolation method to the study of buffers with two binding sites. The feasibility of extending the variational method used in (37) is still an open question and could be addressed in future work. One should explore in particular the applicability of the multifunction variational method described in (53).

More importantly, the newly developed approximants can be used to study in detail the parameter dependence of equilibrium concentrations of Ca^2+^ and distinct buffer states, which can be quite non-trivial for a buffer with two binding sites. For example, the results shown in Figure 1 already reveal an interesting non-monotonic dependence of single-bound buffer on the distance from the Ca^2+^ channel for some, but not all, model parameters. To our knowledge, this non-monotonic behavior has not been previously noted. Since most buffers have dual Ca^2+^ buffering and sensing roles, with partially and fully bound buffer having distinct affinities to downstream biochemical targets (39, 40, 47), this non-trivial property of the equilibrium solution may be of potential physiological significance, to be analyzed in detail. Non-trivial effects of cooperative Ca^2+^ binding by biological buffers with multiple Ca^2+^ binding sites has also been pointed out by prior modeling studies. For example, it has been shown that cooperative Ca^2+^ buffers decrease the facilitation of Ca^2+^ transients associated with buffers saturation (39, 44, 47), but may increase short-term synaptic facilitation through the mechanism of buffer dislocation (56). This is directly related to the interesting fact that the buffering capacity of a cooperative Ca^2+^ buffer increases with increasing background Ca^2+^ concentration, which may play an important homeostatic role (41, 44, 48). On longer time scales, Kubota and Waxham (46) showed the interesting “hand-off” of Ca^2+^ from the N-lobe to the C-lobe of calmodulin upon channel closing, and the intricate dependence of each lobe’s Ca^2+^ saturation on the Ca^2+^ influx amplitude and duration. More generally, cooperative Ca^2+^ binding by calmodulin and the resulting activation of downstream biochemical pathways plays important roles in the regulation of long-term synaptic plasticity and other fundamental cell processes (42-46, 51, 57-59). Deeper understanding of Ca^2+^ dynamics in the presence of cooperative buffers may also be important for an accurate interpretation of optogenetic measurements with genetically-encoded fluorescent Ca^2+^ dyes, which are formed by fusing a calmodulin molecule with a green fluorescent protein (60). All this underscores the importance of modeling and analysis of Ca^2+^ binding by buffers and sensors with multiple Ca^2+^ binding sites.

## Author Contributions

V.M. conceived, designed, and supervised this research project; Y.C. and V.M. both performed model analysis, coding, numerical simulations and analysis of results, and both took part in the writing of the manuscript.

## Acknowledgements

This work was supported in part by NSF grant DMS-1517085 to V.M. The authors acknowledge valuable discussions with Cyrill Muratov and Arthur Sherman.

## A APPENDIX

### Approximation Parameters, Zero Background Ca^2+^ Concentration

1. For **PadéB** and **ExpPadéB** approximation, matching the coefficients of the short- and long-range series expansions given by Eqs. leads to cubic systems for the *ansatz* parameter *b*_2_ shown in Table 3, with the following explicit solution:

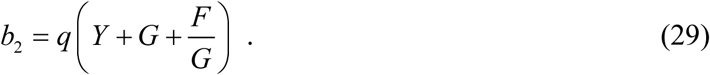
  a. For **PadéB**, the auxiliary quantities *Y, G, F* are determined by

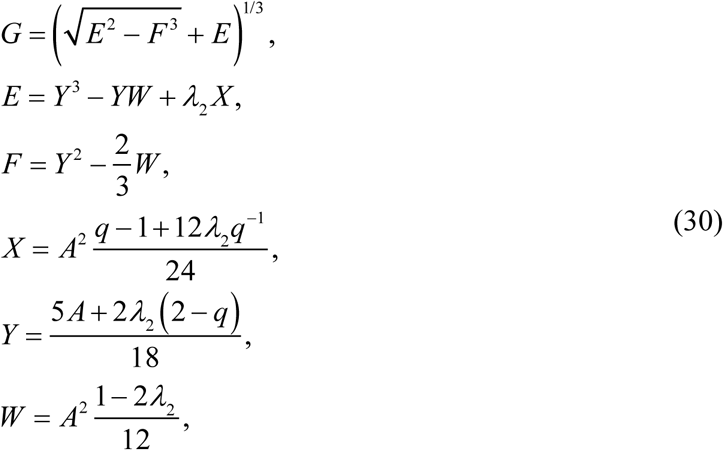

where the value of *ansatz* parameter *A* is shown in Table 2.
  b. For **ExpPadéB**, the computation of *b*_2_ value is the same as above, except for the redefinition of the auxiliary quantity *Y*:

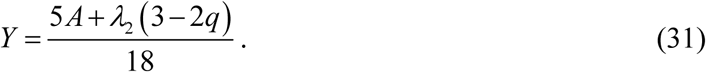
2. For both **PadéExp** and **ExpExp** approximations, the explicit solution of *ansatz* parameter *s* has the same form:

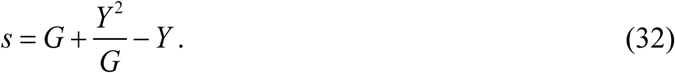

where the auxiliary quantities *G* and *Y* are determined by

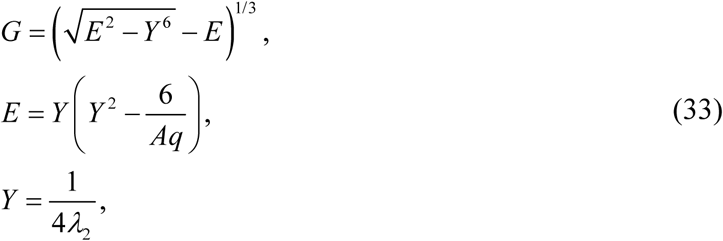

and the value of constant *A* is given in Table 2. □

Supporting Material 2 contains simple code implementing these expressions using MATLAB functions (MathWorks, Inc).

